# A profile-based method for identifying functional divergence of orthologous genes in bacterial genomes

**DOI:** 10.1101/022616

**Authors:** Nicole E. Wheeler, Lars Barquist, Robert A. Kingsley, Paul P. Gardner

**Affiliations:** School of Biological Sciences, University of Canterbury, Christchurch, New Zealand; Biomolecular Interaction Centre, University of Canterbury, Christchurch, New Zealand; Institute for Molecular Infection Biology, University of Wuerzburg, Wuerzburg, Germany; Institute of Food Research, Norwich Research Park, Norwich, UK; Wellcome Trust Sanger Institute, Hinxton, UK; Bio-protection Research Centre, University of Canterbury, Christchurch, New Zealand

## Abstract

**Motivation:** Next generation sequencing technologies have provided us with a wealth of information on genetic variation, but predicting the functional significance of this variation is a difficult task. While many comparative genomics studies have focused on gene flux and large scale changes, relatively little attention has been paid to quantifying the effects of single nucleotide polymorphisms and indels on protein function, particularly in bacterial genomics.

**Results:** We present a hidden Markov model based approach we call delta-bitscore (DBS) for identifying orthologous proteins that have diverged at the amino acid sequence level in a way that is likely to impact biological function. We benchmark this approach with several widely used datasets and apply it to a proof-of-concept study of orthologous proteomes in an investigation of host adaptation in *Salmonella enterica*. We highlight the value of the method in identifying functional divergence of genes, and suggest that this tool may be a better approach than the commonly used dN/dS metric for identifying functionally significant genetic changes occurring in recently diverged organisms.

**Availability:** A program implementing DBS for pairwise genome comparisons is freely available at: https://github.com/UCanCompBio/deltaBS.

**Supplementary information:** Supplementary data are available at BioRxiv online.

## 1 Introduction

Genome sequencing technologies allow us to explore the wealth of genetic variation between and within species, and as these technologies advance this data is becoming progressively cheaper, faster, and easier to produce (Loman et al., 2012; Koren and Phillippy, 2015; Loman and Pallen, 2015). However, analysis of the functional impact of genetic variation has lagged behind, and has largely focused on the presence or absence of macroscopic features such as particular genes, genomic islands, or plasmids. Comparative sequence analyses have become common, and exploration of genetic variation between closely-related organisms has provided key insights into bacterial evolution (Croucher and Didelot, 2014; Bryant et al., 2012; Barquist and Vogel, 2015). In particular, the analysis of single nucleotide polymorphisms (SNPs) has been a tremendous boon to the study of bacterial populations, allowing for the construction of phylogenetic trees which provide information on disease transmission and adaptation at scales ranging from global pandemics (Mutreja et al., 2011) to outbreaks within single hospital wards (Harris et al., 2013). Still, the functional analysis of these SNPs, insertions, and deletions within protein sequences remains difficult, and often relies on inappropriate tools such as dN/dS.

How then can the significance of fine-scale genetic variation be quantified and prioritized for investigation? Recent studies have shown that even single SNPs can have dramatic effects on major phenotypes such as host tropism (Viana et al., 2015; Yue et al., 2015; Singletary et al., 2016). Studies of pathogen adaptation, for example the adaptation of *Pseudomonas aeruginosa* to the cystic fibrosis lung (Marvig et al., 2015; Jorth et al., 2015) or of *Salmonella enterica* to immunocompromised populations (Feasey et al., 2012; Okoro et al., 2012, 2015), often result in findings of hundreds to thousands of SNPs and small indels in coding regions. Genome-wide association studies provide one method (Chewapreecha et al., 2014) for interpreting this variation, however the clonal nature of many pathogens can make such study designs difficult if not impossible to pursue and require large sample sizes to be effective. The development of fast and accurate ways to assess the functional impact of variation between strains and prioritize coding variants for follow-up work is an important step in extracting meaning from comparative analyses.

Our strategy uses a profile HMM-based approach. Profile HMMs are probabilistic models of multiple sequence alignments. For each column in the alignment they capture information on the expected frequency of occurrence of different amino acids, insertions, and deletions. We can then use this information to compute a score, which we call delta-bitscore (DBS) for reasons explained below, that quantifies the divergence of two protein sequences with respect to the conservation patterns captured by the profile HMM. Our key assumption in this is that variation in positions conserved across a protein family are more likely to affect protein function than variation in less conserved positions. Similar assumptions have had success in a number of applications such as the prediction of tRNA pseudogenes (Lowe and Eddy, 1997) and protein folds (Marks et al., 2012). Our approach is unique in both the simplicity of the measure used, and its flexibility, making it well suited to comparative genomic applications.

We demonstrate that our method is effective using several commonly used protein mutagenesis datasets, particularly when used with automatically constructed protein alignments. We provide an example of a comparative genomic analysis, by applying our approach to a whole proteome analysis of a well-studied case of host adaptation in *Salmonella enterica* serovars Enteritidis and Gallinarum, using a purpose built collection of HMMs. We additionally show these results generalize to other host-adapted salmonellae. Finally, we compare our method to a popular estimate of selection, dN/dS, and find that the two measures are only weakly correlated. This adds to the evidence that dN/dS is not an appropriate measure for inferring selection for gene function within bacterial populations.

## 2 Methods

### 3.1 Benchmarking

In order to test the ability of profile HMMs to identify loss-of-function mutations we used three independent datasets from protein mutagenesis studies on HIV-1 protease (336 sequences) (Loeb et al., 1989), E. coli Lac I (4041 sequences) (Markiewicz et al., 1994), and phage lysozyme (2015 sequences) (Rennell et al., 1991). In these experiments, residues in the protein were systematically mutated and the impact of these mutations on protein function was quantified. The data were downloaded from the SIFT website (http://sift.bii.a-star.edu.sg/) (Kumar et al., 2009). For binary classification of mutants, proteins with a classification of “+” were termed functional mutants and “+-”, “-+”, and “-” were termed loss of function mutants. We tested two different reference HMM sets: curated HMMs from the Pfam database (Punta et al., 2012), and automatically constructed HMMs built using a range of residue identity cutoff values. We also compared these results to predictions made by the PROVEAN, SIFT and Mutation Assessor methods. PROVEAN results were downloaded pre-computed from their website. Otherwise, all methods were tested using their default settings. Exclusion of positions in the protein with low coverage in the SIFT database have only a marginal effect on SIFT performance (0.7 to 0.71 AUC in lysozyme dataset), and results in loss of data, so we use all sequence variants in our analysis. In order to evaluate the accuracy of each method we computed the ‘Area Above the Curve’ (AAC) for a series of Relative Cost Curve (RCC) plots (Montvida and Klawonn, 2014). These values range between one and zero with an AAC equal to one indicating a perfect prediction tool across the range of relative costs tested. The Biocomb package (www.cran.rproject.org/package=Biocomb) was used with default relative costs unless otherwise specified to calculate AAC scores, and the ROCR package (Sing et al., 2005) was used to calculate other performance metrics, which can be found in Supplementary Table 1.

For the construction of custom HMMs, query sequences were searched against the Uniref90 database (Suzek et al., 2015) using a single iteration of jackhmmer. Pairwise sequence identity was calculated as the number of matches between sequence pairs, divided by total length of the alignment after the removal of gap-gap columns. HMMs were built using hmmbuild, and DBS was calculated using hmmsearch, subtracting the bitscore value for the variant sequence from the bitscore value for the wild type sequence. jackhmmer, hmmbuild and hmmsearch are part of the HMMER3 package (version 3.1b2). Protein HMMs were built using a range of sequence identity cutoffs in order to determine the optimal range for performance.

### 3.2 Proteome analysis

We have designed this approach with whole proteome analyses in mind, so as a proof-of-concept, we applied the approach to the study of host adaptation in *Salmonella enterica*, using *S*. Enteritidis and *S*. Gallinarum as a specific example. For a summary of the workflow of this portion of the analysis, see Supplementary Figure 1. Genomes for *Salmonella* Enteritidis str. P125109 (AM933172.1) and *Salmonella* Gallinarum str. 287/91 (AM933173.1) were retrieved. Custom gene models were constructed by searching each S. Enteritidis protein coding gene against the Uniref90 database. Profiles were built using sequences showing 40% or greater sequence identity to the original query protein.

**Fig. 1.**
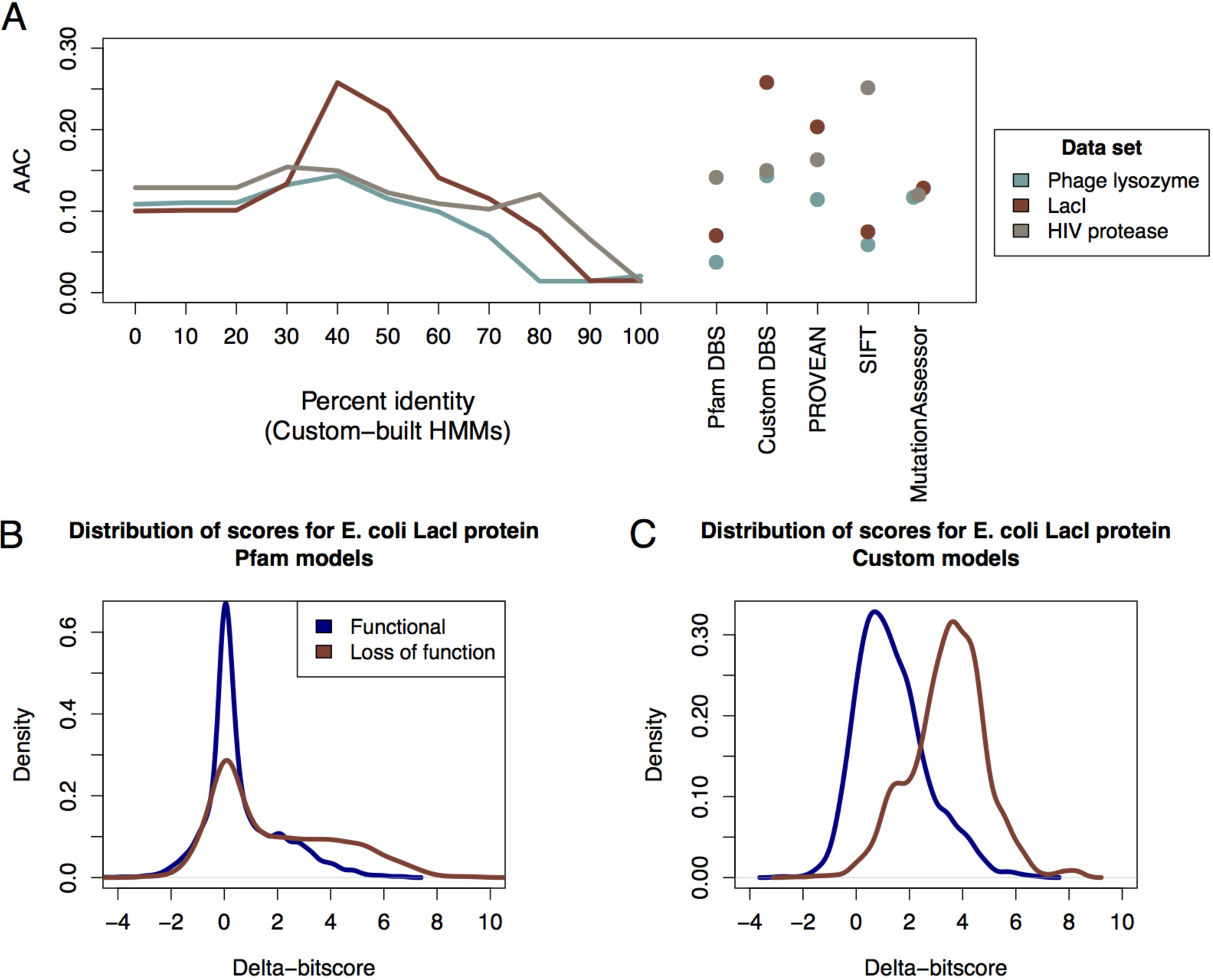
**(A)** Area above the relative cost curve (AAC) values for different analysis methods using three mutagenesis benchmarking datasets: phage lysozyme, LacI and HIV protease. The performance for each model across a range of sequence identity cutoffs is shown, as well as AAC values for the individual methods. AAC of 1.0 indicates perfect performance.; **(B)** Distribution of delta-bitscores for *E. coli* LacI using custom model analysis; **(C)** Distribution of delta-bitscores for *E. coli* LacI using Pfam model analysis.

In order to assess whether models built from a small number of sequences performed well, we tested the performance of models using proteins from the humsavar database (http://www.uniprot.org/docs/humsavar), which catalogs human polymorphisms and disease variants for a wide range of proteins which have different levels of representation in Uniprot90, and therefore result in profile models built from varying numbers of sequences. We built custom models for each protein in the database, then separated the models into classes based on both number of sequences and effective sequence number. We then computed AUC values for predictions made on variant data from each class. Results from this test can be found in Supplementary Table 2. We saw no classes where custom models performed worse than Pfam models, so we did not filter models built from few sequences. We did, however, use Pfam models for those proteins with no homologs in Uniref90 rather than build a model from the query sequence, as we felt this could bias results towards the reference species.

To assess the quality of our analysis, we wished to compare our results to those of Nuccio and Bäumler (2014), so we used the ortholog calls provided in their supplementary material. Genes represented by a custom model were scored using the appropriate model (n=3154), all others were scored using Pfam domain models. hmmsearch was used for scoring. If hits to multiple Pfam models occurred, any overlapping hits were competed based on E-value. Orthologs with incompatible Pfam domain architectures (n=32) were excluded from scoring, but counted as hypothetically attenuated coding sequences (HACs) in Figure 2A if they involved a loss of domain in one serovar. We anticipate that the variance of delta-bitscores will increase with evolutionary distance, so rather than establish a fixed scoring cutoff for HACs, we identify loss-of-function mutations using an empirical distribution. We set a score threshold at which 2.5% of genes on the least dispersed side of the distribution would be called as HACs. If the two bacteria show an equal rate of protein function loss, this would result in 5% of orthologous proteins being called as HACs, however if one proteome shows a greater degree of functional degradation, this will result in a greater proportion of orthologs being classified as HACs.

**Fig. 2.**
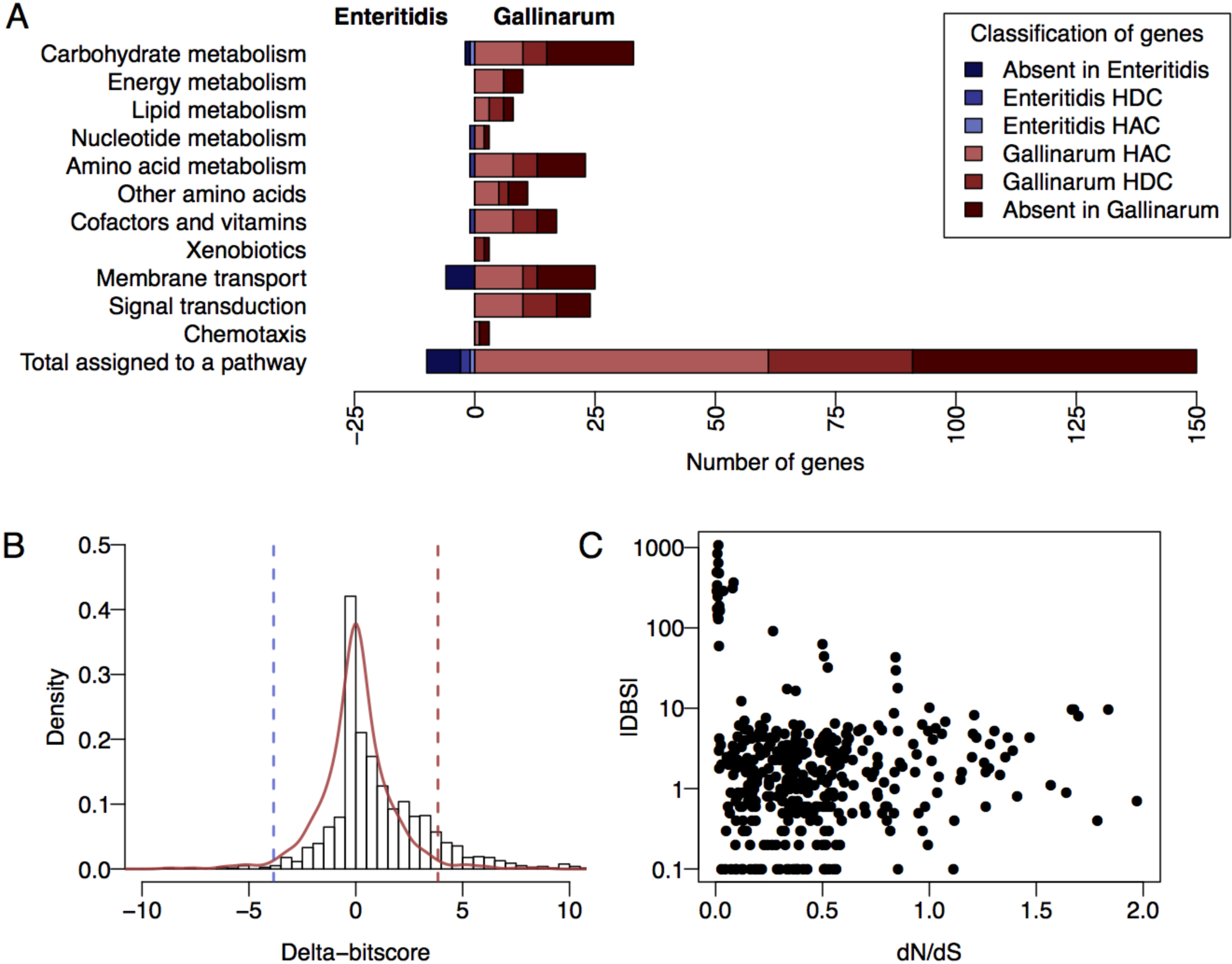
The results of a DBS comparison of the proteomes of *S.* Enteritidis (a generalist infectious agent) and *S.* Gallinarum (a host-restricted infectious agent). **(A)** The functional changes in orthologous protein-coding genes of *S.* Enteritidis and *S.* Gallinarum have been grouped into functional categories using the KEGG pathways database. Genes included in the pathway that have no ortholog in the other serovar are indicated in the darkest colour, followed by genes previously identified as hypothetically disrupted coding sequences (HDCs), then genes with significant DBS values that had not already been identified as HDCs. We refer to these as hypothetically attenuated coding sequences (HACs); **(B)** The distribution of delta-bitscores for orthologous genes showing non-synonymous changes in *S.* Enteritidis and *S.* Gallinarum. A symmetrical empirical distribution of scores generated by mirroring the least dispersed side is shown in red, the cutoff values we use to establish significance are shown as dashed lines. A skewed distribution implies excess functional changes in one lineage; **(C)** A plot of the dN/dS score and corresponding DBS is shown for each orthologous *S.* Enteritidis and *S.* Gallinarum gene.

Functional classification was performed using pathways from the KEGG database (Kanehisa et al., 2016). We grouped genes into four categories: those present in a pathway but with no ortholog in the other serovar; genes identified as hypothetically disrupted coding sequences (HDCs) by Nuccio and Bäumler (2014); genes identified by our DBS method as HACs, but not as HDCs; and finally genes not identified as nonfunctional by either method. dN/dS values were calculated using PAML (Yang, 1997), and for the comparison of DBS and dN/dS, genes were filtered for those with dN>0 and dS>0.0001. Correlations between measures were computed using a Spearman’s rho statistic (R package cor).

In our investigation of multiple *S. enterica* isolates, for all orthologous groups with a gene present in *S*. Enteritidis, scores were collated, and if individual scores were significantly different to the median score for all isolates, we identified these proteins as HACs. Significant difference was calculated in a similar way to the pairwise comparisons, with the score corresponding to the most extreme 2.5% of delta-bitscores on the least dispersed side of an empirical distribution being used as the cutoff.

## 3 Results

### 3.3 The delta-bitscore measure

Alignment of a protein sequence to a profile HMM produces a bitscore value, which is the log odds ratio for this sequence under the profile HMM compared to a random null model, and serves as an indication of protein family membership. In subtracting the bitscore of one protein or domain from that of another, we produce a measure of the divergence between the two proteins, taking into consideration conservation patterns captured by the protein family model.

We define delta-bitscore using the following equation:

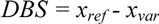

Where DBS is delta-bitscore and, *x_ref_* and *x_var_* are bitscores for reference and variant sequences derived from alignments to the same profile HMM.

Highly conserved positions in a model alignment contribute more to bitscores than poorly conserved positions, meaning that unexpected mutations or indels in conserved sites are given a greater penalty than mutations in variable sites. In addition, the replacement of residues with chemically and structurally similar residues is generally scored more favourably than mutation to dissimilar residues. Multiple functionally neutral changes are likely to result in individual contributions to the bitscore that are small in magnitude and that cancel out over the length of the protein, while functionally significant change in a protein will likely produce one or more position-specific values of high magnitude that have a greater impact on overall DBS for the sequence. We first introduced this measure as part of studies of *Salmonella* adaptation to an wild avian host (Kingsley et al., 2013), and during within-host evolution of a hypermutator strain of *Salmonella* Enteritidis in an immunocompromised patient (Klemm et al., 2016), and elaborate upon it here. The Supplementary Text contains further discussion of this measure and comparisons with related measures derived from profile HMMs (Clifford et al., 2004; Shihab et al., 2013; Liu et al., 2015).

### 3.4 DBS is predictive of protein functional status

Evaluating the performance of different methods for the quantification of the effects of sequence variation is challenging. For the case of human variation, some datasets exist, however given the limited amount of data available on functional protein variants care must be taken to avoid circularity in training and testing (Boulesteix, 2010; Grimm et al., 2015). While we have not comprehensively benchmarked DBS on human data sets given our focus on prokaryotic variation, we found it is competitive with other untrained methods (Supplementary Figures 3, 4), despite its simplicity. In the case of bacterial variation we are not aware of well-characterized collections of protein variants. Rather, to test our method we compared its performance to a selection of methods on three large scale protein mutagenesis datasets (Kumar et al., 2009) (Figure 1A, Supplementary Figure 2 shows AUC values for the same analysis).

A number of methods have been designed for predicting the impact of sequence variation on the function of non-human proteins, and we compare our measure to three competitive methods here (Grimm et al., 2015). These include: PROVEAN, which uses a BLAST-based approach to score sequence variants using closely related sequences (Choi et al., 2012); the SIFT algorithm which uses position-specific scoring matrices based on sequence homology and known patterns of common amino acid substitutions to predict the functional consequences of non-synonymous single nucleotide polymorphisms (nsSNPs) (Kumar et al., 2009); and MutationAssessor which computes both the conservation and specificity of residues within protein subfamilies to assess the impact of a mutation (Reva et al., 2011).

As a comparison, we used DBS in two distinct ways. In the first, we calculated DBS scores based on alignments to curated Pfam domains (Punta et al., 2012). In the second, we constructed profile-HMMs containing sequences of varying minimal sequence identity. These two approaches to model construction capture distinct forms of information: Pfam domains are designed to capture maximal sequence diversity, and focus on deeply conserved domains within protein sequences. Additionally, many families are manually curated, and the alignment of known functional regions, such as active sites, may have been improved in some cases. Our custom profiles capture full length protein sequences. This means that the functions of sequences captured by custom models may be narrower than those captured by Pfam, and additionally will capture linker regions between conserved domains, which may nevertheless be functionally important.

Our benchmark showed that custom HMMs built at 40% identity outperformed other methods in terms of area above the relative cost curve (AAC) and maximum Matthew’s correlation coefficient, with the exception of the superior performance of SIFT and PROVEAN on the HIV protease data (Figure 1A, and Supplementary Table 1). DBS using Pfam models had the best specificity across all datasets tested, but at the expense of sensitivity (see Figure 1B, Supplementary Table 1). This suggests that Pfam models are indeed capturing ancient features of protein families, and may be useful in cases where a low false-positive rate is desired at the expense of potentially missing variants. Our custom models meanwhile appeared to perform optimally with a 40% identity cutoff (Figure 1A, C). Interestingly, 40% sequence identity has previously been proposed as a rule of thumb identity cutoff for the transfer of enzyme annotations (Addou et al., 2009; Tian and Skolnick, 2003), indicating that this cutoff may reflect a common point of functional divergence in protein families.

### 3.5 DBS identifies pathways associated with host range restriction in Salmonella Gallinarum

We have established that DBS can detect functionally relevant mutations. To explore the utility of this approach in a comparative genomics context, we developed a tool that takes whole proteome files as input, builds custom HMMs where applicable and uses Pfam HMMs to score the remaining genes, in order to identify functionally significant variation between two proteomes. Using this tool, we compared the proteomes of two closely related *Salmonella enterica* serovars that have experienced contrasting selective pressures leading to host-restriction for one of them.

Host-restriction is a common phenomenon in highly adapted invasive pathogens, often characterized by genomic features such as the proliferation of transposable elements and the degradation of substantial fractions of coding sequences (Moran and Plague, 2004; Goodhead and Darby, 2015). Within *Salmonella enterica* such restriction events have occurred independently multiple times in various hosts from broad host-range ancestors. *Salmonella enterica* serovar Enteritidis is a broad host-range pathogen, capable of infecting humans, cattle, rodents and a variety of birds, while the closely related serovar Gallinarum is restricted to infecting galliforme birds (Rabsch et al., 2002). *S.* Gallinarum and *S.* Enteritidis have recently evolved from a common ancestor, however the *S*. Gallinarum genome has undergone extensive degradation since divergence (Thomson et al., 2008; Langridge et al., 2015). In addition to being restricted to a narrow host range, *S.* Gallinarum has lost motility and causes a systemic, typhoid-like infection in birds, unlike Enteritidis which usually causes a self-limiting gastroenteritis (Thomson et al., 2008). A recent analysis of pseudogenes within this and other invasive *Salmonella* lineages identified signatures of host-restriction, characterized particularly by the loss of metabolic genes required for survival in the intestine (Langridge et al., 2015; Nuccio and Bäumler, 2014). Both of these analyses involved the manual comparison of coding sequences to identify mutations that would result in frameshifts or truncated proteins. We expect DBS to add an additional layer of information to such an analysis, identifying genes which have shifted in or lost function due to non-synonymous mutations or small indels occurring since the restriction event, but have not yet succumbed to obvious disruption events such as large truncations, frameshifts, or complete deletions.

As shown in Figure 2A, our loss-of-function predictions included many genes not identified as hypothetically disrupted coding sequences (HDCs) by manual inspection (Nuccio and Bäumler, 2014).

We examined the concurrence between our loss of function calls and those made by Nuccio and Bäumler. Of 252 HDCs identified in *S.* Gallinarum but not in *S.* Enteritidis from the previous study, 148 of these were also classified as HACs by DBS, and a further 6 as HACs due to a loss of a domain in the *S.* Gallinarum copy of the gene (61% total agreement). Of the 104 disagreements, 28 genes had a DBS score below the threshold, and 70 were excluded from scoring analysis for having incompatible domain architectures (n=14) or no hits for either protein sequence to the Pfam database or any of our custom models (n=56). Of these 56, over half were fewer than 150 amino acids long, often due to extreme truncations or early frameshifts (see Supplementary Table 3). Of the 28 low scoring variants, many involved short truncations, alternate starts or indels of <10 amino acids. The remainder involved *S.* Gallinarum pseudogenes that had no hits to custom models or Pfam domains in the truncated region. In many cases our models showed low sequence conservation and gaps in the regions affected by mutation.

As expected, the distribution of DBS values centers around zero (Figure 2B), showing that most orthologous gene pairs differ from the modeled sequence constraints to a similar degree. The distribution of DBS values shows an enrichment for positive DBS (exact binomial test, P = 2.18 e-35), indicating greater divergence from the profile models for protein coding genes from *S.* Gallinarum when compared to *S.* Enteritidis. While some of these positive DBS values may indicate functional divergence rather than pseudogenization, extreme DBS values predominantly correspond to truncations, indels and mutations in highly conserved sites, so we assume that in these cases the ancestral function has been lost. To look at these hypothetically attenuated coding sequences (HACs) in a functional context, we grouped genes into functional categories based on their annotation in the KEGG database. Not only does *S.* Gallinarum have fewer genes than *S*. Enteritidis for most of the functional categories we considered (data not shown), but it also has a greater number of HACs across these groupings (Figure 2A). Previous work that found the presence of non-ancestral pseudogenes in *S.* Enteritidis was limited while *S.* Gallinarum had accumulated a large number of pseudogenes since divergence is consistent with our results (Langridge et al., 2015).

In our analysis we found a number of results which are in agreement with previously published findings. Degradation of genes involved in chemotaxis is consistent with a loss of motility in this strain, and previously identified degradation of chemotaxis genes in other host adapted strains (Kingsley et al., 2013; McClelland et al., 2004). Fimbriae are also thought to be important for host colonisation by *Salmonella*, and we found high DBS values for bcfA, bcfC and stfG. Pseudogenization of other genes in these operons have already been identified in *S.* Gallinarum (Foley et al., 2013). We found a number of HACs in genes involved in the utilization of nutrients derived from the inflamed host gut environment colonized by gastrointestinal serovars of *Salmonella* (Nuccio and Bäumler, 2014) (Supplementary Table 5). This is consistent with the recent adaptation of *S.* Gallinarum from an ancestral gastrointestinal pathogen to an extra-intestinal environment. Among the most highly ranked genes were representatives of the cbi, pdu, and eut operons, previously identified as central pathways subject to gene decay in *S.* Gallinarum (Langridge et al., 2015; Nuccio and Bäumler, 2014). A complete table of DBS values for orthologous genes can be found in Supplementary Table 6.

These findings demonstrate that DBS provides an additional layer of information about gene function in addition to gene deletion when investigating serovars that have recently diverged. As time since divergence increases we expect that non-functional genes will be deleted and the ratio of non-functional and deleted genes will change. We anticipate that our method will be most useful in comparisons of organisms which have recently diverged, as it offers an opportunity to identify loss-of-function mutations that occur as an immediate response to a new environment and restricted population size, before deletion of entire genes occurs (Kuo and Ochman, 2010).

### 3.6 DBS does not correlate strongly with dN/dS

In comparing genome sequences, we have focused on identifying functionally significant variation, whereas a common approach is to classify genes from an evolutionary point of view, i.e. to identify genes are under negative selection, positive selection or that are evolving neutrally. A commonly used measure of selection is dN/dS (ω), which compares the rate of nonsynonymous changes in protein-coding sequences to the rate of synonymous changes to classify genes as either negative selection (ω < 1), positive selection (ω > 1) or neutrally evolving (ω ≈ 1) (Yang and Bielawski, 2000). While dN/dS is an inappropriate metric to use in comparing bacteria at the strain level (Rocha et al., 2006), the metric is nevertheless used in similar situations (Roumagnac et al., 2006; Fleischmann et al., 2002; Holden et al., 2004), so we compared the results of the commonly used PAML program and DBS. In comparing the results from the two approaches, one of the most striking results is that there is little correlation between DBS and both dN/dS (R ≈ 0.17) and dN (R ≈ 0.18). Of the genes with high DBS and low dN/dS, most of these can be attributed to insertions and deletions, which are ignored by dN/dS but can have significant impacts on the functioning of a protein. Genes with low DBS and high dN/dS tend to carry a small number of nonsynonymous changes, most of which are between chemically similar residues, and a smaller number of synonymous mutations.

### 3.7 DBS reveals trends of gene degradation across host-adapted Salmonella

To test whether a positively skewed DBS distribution was a common feature of host adaptation in *Salmonella*, we performed DBS analysis on three broad host range and three host-adapted *Salmonella* serovars. Rather than performing many pairwise comparisons to call HACs, for this analysis we compared bitscores for each gene to the median gene bitscore of the six strains. We found that host-restricted serovars showed a greater number of HACs than generalist serovars across all strains tested (Figure 3A). We previously observed a similar phenomenon in host-restricted strains of *S.* Typhimurium (Kingsley et al., 2013). We observed a striking similarity in the score distributions of generalist strains, and in host-adapted strains (Supplementary Figure 6), with the generalist scores being lower overall than the host restricted scores. It is interesting to note that most of the separation in scores occurs at DBS values below our cutoff value for HAC classification (DBS = 8.2, Supplementary Figure 6, top panel). Based on our benchmarking, if these proteins were tested in vitro many of them would show differences in function, but whether these differences relate to differences in intracellular and environmental conditions shaping the sequence requirements for these proteins or whether they relate to true declines in function is difficult to assess without experimental validation.

**Fig. 3.**
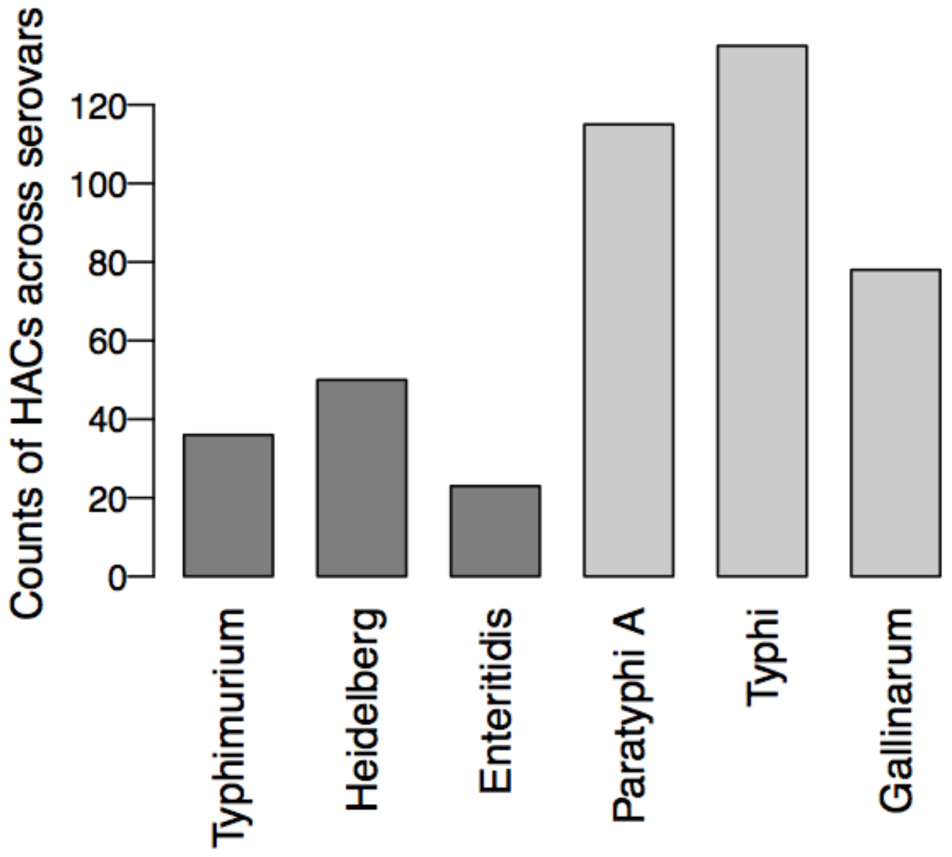
Counts of hypothetically attenuated coding sequences (HACs) for each serovar, identified as the most extreme deviations from the median bitscore across strains for each gene. The generalist serovars are coloured in dark grey, host-restricted serovars are coloured in light grey.

## 4 Discussion

We have described a surprisingly simple method for identifying functional divergence in orthologous proteins, including analysis of both point mutations and indel events. Despite the simplicity of the DBS measure, its accuracy is similar to state-of-the-art methods for predicting the functional impact of sequence variants. While curated Pfam models were less sensitive, they have a high specificity at conservative scoring thresholds (see Supplementary Table 1, Supplementary Figure 5). The workflow for using such models is simpler (see Supplementary Figure 1), and may be more appropriate for investigations where a low false-positive rate is more important than detection of all loss-of-function mutations. In addition, while construction of custom models can be time-consuming to run on a desktop computer, Pfam annotation can be performed rapidly. Because other methods for estimating the impact of sequence variation, such as SIFT, PROVEAN and MutationAssessor, require the user to specify mutations individually, broad scale comparative proteome analysis such as this is difficult and time-intensive to perform, and not all methods are able to score indels or more than one mutation in the same sequence. Both DBS approaches scale to whole proteome analysis with minimal user involvement, and were able to detect additional loss of function mutations that were missed by previous studies focused on protein truncations and frameshifts.

### 3.8 DBS for the study of bacterial adaptation

Large scale genomic studies of bacterial pathogens have revealed striking similarities in evolutionary patterns in diverse pathogenic and symbiotic lineages. While adaptation to a new environment may involve the acquisition of new fitness determinants and rare, beneficial mutations, niche adaptation is also frequently accompanied by widespread loss-of-function mutations, particularly in pathogenic and symbiotic lineages. These loss-of-function mutations are likely generated through a number of distinct processes (Moran, 2002), including neutral or slightly deleterious stochastic loss due to reductions in effective population size, neutral loss of genes no longer required in the new environment, and adaptive loss (Hottes et al., 2013). DBS presents an opportunity to mine the genomes of bacteria adapting to new environments for these loss of function mutations, which may tell us as much about bacteria’s adaptation to new environments as the acquisition of new genes. Comparative approaches based on pseudogene analysis have identified consistent signatures of adaptation in a number of bacterial pathogens, including *Yersinia* (Reuter et al., 2014; McNally et al., 2016), *E. coli* and *Shigella* (Monk et al., 2013; Feng et al., 2011), and *Salmonella* (Langridge et al., 2015; Nuccio and Bäumler, 2014). Our analysis of *Salmonella* genomes has shown that DBS is consistent with these previous studies, while providing additional sensitivity to detect non-functional or functionally divergent protein variants that may have been missed by pseudogene analyses (Kingsley et al., 2013).

### 3.9 DBS as a more appropriate alternative to dN/dS for studying evolution across short time scales

While dN/dS can be a powerful approach to identifying genes under positive selection across long time scales, on shorter timescales it has been shown to be inaccurate, due to the lag in the removal of slightly deleterious mutations This leads to high dN/dS ratios being commonplace in comparisons of closely related strains, suggesting positive or relaxed selection where there is none (Rocha et al., 2006). Other studies have shown that dN/dS can provide inaccurate, even contradictory, results depending on the structure of the population being sampled (Kryazhimskiy and Plotkin, 2008). Due to the unreliability of dN/dS measures at short evolutionary timescales and the inability of dN/dS based methods to score indels, we propose that DBS is a more suitable analysis tool for the study of functional divergence in recently diverged strains. In contrast, we would advise caution in the use of DBS on more divergent organisms, with careful consideration of the effects of sequence divergence on bitscore. Over time each lineage could accumulate mutations that affect protein function in different ways, but have the same degree of impact on bitscore, leading to a net DBS near zero. This does not necessarily indicate preservation of function and may instead indicate equal divergence from ancestral function. To date, we have not examined the performance of DBS on deeply diverged lineages. DBS performs optimally when reference alignments are in the 40 percent.identity range, suggesting that sequences more divergent than this may be outside the range where can DBS provide meaningful information, but this will require additional investigation to establish.

### 3.10 Concluding remark

We anticipate that this approach will be a useful addition to traditional comparative genomics workflows. It provides an effective method for scoring the functional potential of the overwhelming numbers of non-synonymous variants that can distinguish closely related bacteria.

## Acknowledgements

We would like to thank Fatemeh Ashari Ghomi for her contributions, and Sean Eddy for his clarification of some technical aspects of the HMMER3 software.

## References

Addou, S. et al. (2009) Domain-based and family-specific sequence identity thresholds increase the levels of reliable protein function transfer. J. Mol. Biol., 387, 416–430.

Barquist, L. et al. (2013) Approaches to querying bacterial genomes with transposon-insertion sequencing. RNA Biol., 10, 1161–1169.

Barquist, L. and Vogel, J. (2015) Accelerating Discovery and Functional Analysis of Small RNAs with New Technologies. Annu. Rev. Genet., 49, 367–394.

Boulesteix, A.-L. (2010) Over-optimism in bioinformatics research. Bioinformatics, 26, 437–439.

Bryant, J. et al. (2012) Developing insights into the mechanisms of evolution of bacterial pathogens from whole-genome sequences. Future Microbiol., 7, 1283–1296.

Chewapreecha, C. et al. (2014) Comprehensive identification of single nucleotide polymorphisms associated with beta-lactam resistance within pneumococcal mosaic genes. PLoS Genet., 10, e1004547.

Choi, Y. et al. (2012) Predicting the functional effect of amino acid substitutions and indels. PLoS One, 7, e46688.

Clifford, R.J. et al. (2004) Large-scale analysis of non-synonymous coding region single nucleotide polymorphisms. Bioinformatics, 20, 1006–1014.

Croucher, N.J. and Didelot, X. (2014) The application of genomics to tracing bacterial pathogen transmission. Curr. Opin. Microbiol., 23C, 62–67.

Eddy, S.R. (2011) Accelerated Profile HMM Searches. PLoS Comput. Biol., 7, e1002195.

Feasey, N.A. et al. (2012) Invasive non-typhoidal salmonella disease: an emerging and neglected tropical disease in Africa. Lancet, 379, 2489–2499.

Feng, Y. et al. (2011) Gene decay in Shigella as an incipient stage of host-adaptation. PLoS One, 6, e27754.

Fleischmann, R.D. et al. (2002) Whole-Genome Comparison of Mycobacterium tuberculosis Clinical and Laboratory Strains. J. Bacteriol., 184, 5479–5490.

Foley, S.L. et al. (2013) Salmonella pathogenicity and host adaptation in chicken-associated serovars. Microbiol. Mol. Biol. Rev., 77, 582–607.

Goodhead, I. and Darby, A.C. (2015) Taking the pseudo out of pseudogenes. Curr. Opin. Microbiol., 23C, 102–109.

Grimm, D.G. et al. (2015) The evaluation of tools used to predict the impact of missense variants is hindered by two types of circularity. Hum. Mutat., 36, 513–523.

Harris, S.R. et al. (2013) Whole-genome sequencing for analysis of an outbreak of meticillin-resistant Staphylococcus aureus: a descriptive study. Lancet Infect. Dis., 13, 130–136.

Holden, M.T.G. et al. (2004) Complete genomes of two clinical Staphylococcus aureus strains: evidence for the rapid evolution of virulence and drug resistance. Proc. Natl. Acad. Sci. U. S. A., 101, 9786–9791.

Hottes, A.K. et al. (2013) Bacterial adaptation through loss of function. PLoS Genet., 9, e1003617.

Jorth, P. et al. (2015) Regional Isolation Drives Bacterial Diversification within Cystic Fibrosis Lungs. Cell Host Microbe, 18, 307–319.

Kanehisa, M. et al. (2016) KEGG as a reference resource for gene and protein annotation. Nucleic Acids Res., 44, D457–62.

Kingsley, R.A. et al. (2013) Genome and transcriptome adaptation accompanying emergence of the definitive type 2 host-restricted Salmonella enterica serovar Typhimurium pathovar. MBio, 4, e00565–13.

Klemm, E.J. et al. (2016) Emergence of host-adapted Salmonella Enteritidis through rapid evolution in an immunocompromised host. Nature Microbiology, 1, 15023.

Koren, S. and Phillippy, A.M. (2015) One chromosome, one contig: complete microbial genomes from long-read sequencing and assembly. Curr. Opin. Microbiol., 23C, 110–120.

Kryazhimskiy, S. and Plotkin, J.B. (2008) The population genetics of dN/dS. PLoS Genet.

Kumar, P. et al. (2009) Predicting the effects of coding non-synonymous variants on protein function using the SIFT algorithm. Nat. Protoc., 4, 1073–1081.

Kuo, C.-H. and Ochman, H. (2010) The extinction dynamics of bacterial pseudogenes. PLoS Genet., 6.

Langridge, G.C. et al. (2015) Patterns of genome evolution that have accompanied host adaptation in Salmonella. Proc. Natl. Acad. Sci. U. S. A., 112, 863–868.

Liu, M. et al. (2015) HMMvar-func: a new method for predicting the functional outcome of genetic variants. BMC Bioinformatics, 16, 351.

Loeb, D.D. et al. (1989) Complete mutagenesis of the HIV-1 protease. Nature, 340, 397–400.

Loman, N.J. et al. (2012) High-throughput bacterial genome sequencing: an embarrassment of choice, a world of opportunity. Nat. Rev. Microbiol., 10, 599–606.

Loman, N.J. and Pallen, M.J. (2015) Twenty years of bacterial genome sequencing. Nat. Rev. Microbiol., 13, 787–794.

Lowe, T.M. and Eddy, S.R. (1997) tRNAscan-SE: a program for improved detection of transfer RNA genes in genomic sequence. Nucleic Acids Res., 25, 955–964.

Markiewicz, P. et al. (1994) Genetic studies of the lac repressor. XIV. Analysis of 4000 altered Escherichia coli lac repressors reveals essential and non-essential residues, as well as ‘spacers’ which do not require a specific sequence. J. Mol. Biol., 240, 421–433.

Marks, D.S. et al. (2012) Protein structure prediction from sequence variation. Nat. Biotechnol., 30, 1072–1080.

Marvig, R.L. et al. (2015) Convergent evolution and adaptation of Pseudomonas aeruginosa within patients with cystic fibrosis. Nat. Genet., 47, 57–64.

McClelland, M. et al. (2004) Comparison of genome degradation in Paratyphi A and Typhi, human-restricted serovars of Salmonella enterica that cause typhoid. Nat. Genet., 36, 1268–1274.

McNally, A. et al. (2016) ‘Add, stir and reduce’: Yersinia spp. as model bacteria for pathogen evolution. Nat. Rev. Microbiol., 14, 177–190.

Monk, J.M. et al. (2013) Genome-scale metabolic reconstructions of multiple Escherichia coli strains highlight strain-specific adaptations to nutritional environments. Proc. Natl. Acad. Sci. U. S. A., 110, 20338–20343.

Montvida, O and Klawonn, K. (2014) Relative cost curves: An alternative to AUC and an extension to 3-class problems. Kybernetika, 50, 647–660.

Moran, N.A. (2002) Microbial minimalism: genome reduction in bacterial pathogens. Cell, 108, 583–586.

Moran, N.A. and Plague, G.R. (2004) Genomic changes following host restriction in bacteria. Curr. Opin. Genet. Dev., 14, 627–633.

Mutreja, A. et al. (2011) Evidence for several waves of global transmission in the seventh cholera pandemic. Nature, 477, 462–465.

Nozawa, M. et al. (2009) Reliabilities of identifying positive selection by the branch-site and the site-prediction methods. Proc. Natl. Acad. Sci. U. S. A., 106, 6700–6705.

Nuccio, S.-P. and Bäumler, A.J. (2014) Comparative analysis of Salmonella genomes identifies a metabolic network for escalating growth in the inflamed gut. MBio, 5, e00929–14.

Okoro, C.K. et al. (2012) Intracontinental spread of human invasive Salmonella Typhimurium pathovariants in sub-Saharan Africa. Nat. Genet., 44, 1215–1221.

Okoro, C.K. et al. (2015) Signatures of adaptation in human invasive Salmonella Typhimurium ST313 populations from sub-Saharan Africa. PLoS Negl. Trop. Dis., 9, e0003611.

Punta, M. et al. (2012) The Pfam protein families database. Nucleic Acids Res., 40, D290–301.

Rabsch, W. et al. (2002) Salmonella enterica serotype Typhimurium and its host-adapted variants. Infect. Immun., 70, 2249–2255.

Rennell, D. et al. (1991) Systematic mutation of bacteriophage T4 lysozyme. J. Mol. Biol., 222, 67–88.

Reuter, S. et al. (2014) Parallel independent evolution of pathogenicity within the genus Yersinia. Proc. Natl. Acad. Sci. U. S. A., 111, 6768–6773.

Reva, B. et al. (2011) Predicting the functional impact of protein mutations: application to cancer genomics. Nucleic Acids Res., 39, e118.

Rivera-Chávez, F. et al. (2013) Salmonella uses energy taxis to benefit from intestinal inflammation. PLoS Pathog., 9, e1003267.

Rocha, E.P.C. et al. (2006) Comparisons of dN/dS are time dependent for closely related bacterial genomes. J. Theor. Biol., 239, 226–235.

Roumagnac, P. et al. (2006) Evolutionary history of Salmonella typhi. Science, 314, 1301–1304.

Shihab, H.A. et al. (2013) Predicting the functional, molecular, and phenotypic consequences of amino acid substitutions using hidden Markov models. Hum. Mutat., 34, 57–65.

Singletary, L.A. et al. (2016) Loss of Multicellular Behavior in Epidemic African Nontyphoidal Salmonella enterica Serovar Typhimurium ST313 Strain D23580. MBio, 7.

Sing, T. et al. (2005) ROCR: visualizing classifier performance in R. Bioinformatics, 21, 3940–3941.

Suzek, B.E. et al. (2015) UniRef clusters: a comprehensive and scalable alternative for improving sequence similarity searches. Bioinformatics, 31, 926–932.

Thomson, N.R. et al. (2008) Comparative genome analysis of Salmonella Enteritidis PT4 and Salmonella Gallinarum 287/91 provides insights into evolutionary and host adaptation pathways. Genome Res., 18, 1624–1637.

Tian, W. and Skolnick, J. (2003) How well is enzyme function conserved as a function of pairwise sequence identity? J. Mol. Biol., 333, 863–882.

Viana, D. et al. (2015) A single natural nucleotide mutation alters bacterial pathogen host tropism. Nat. Genet.

Yang and Bielawski (2000) Statistical methods for detecting molecular adaptation. Trends Ecol. Evol., 15, 496–503.

Yang, H. et al. (2014) Genome-Scale Metabolic Network Validation of Shewanella oneidensis Using Transposon Insertion Frequency Analysis. PLoS Comput. Biol., 10, e1003848.

Yang, Z. (1997) PAML: a program package for phylogenetic analysis by maximum likelihood. Comput. Appl. Biosci., 13, 555–556.

Yue, M. et al. (2015) Allelic variation contributes to bacterial host specificity. Nat. Commun., 6, 8754.

